## Supplementary Tables S1-S7 for "Maintenance of homeostatic plasticity at the *Drosophila* neuromuscular synapse requires continuous IP_3_-directed signaling"

**Supplementary Tables S1-S7. Summary Electrophysiological Data.** Genotypes and rearing conditions for electrophysiological data presented in the study (corresponding figures specified). For GAL4 drivers, "*Pre + Post-Gal4*" denotes a genetic combination of *elaV(C155)-Gal4/Y; Sca-Gal4/+; BG57-Gal4/+* (see Materials and Methods). Average values ± SEM are presented for each electrophysiological parameter, with *n* = number of NMJs recorded. Values include miniature excitatory postsynaptic potential (mEPSP) amplitude, mEPSP frequency (Freq), excitatory postsynaptic potential (EPSP) amplitude, quantal content (QC), and QC corrected for non-linear summation (NLS). * *p* < 0.05, ** *p* < 0.01, *** *p* < 0.001 by Student's T-Test vs. unchallenged control.

### Supplementary Table S1

| **FIGURE 1** | | | | | | | | |
|---|---|---|---|---|---|---|---|---|
| **Condition** | **Genotype or Reagent** | **mEPSP (mV)** | **mEPSP freq. (Hz)** | **EPSP (mV)** | **$V_m$ (mV)** | **QC** | **NLSC QC** | **n** |
| Driver control | *Pre + Post-Gal4* | 0.83 ± 0.03 | 1.0 ± 0.1 | 40.4 ± 1.4 | -69.0 ± 1.2 | 48.6 ± 1.1 | 100.2 ± 3.7 | 13 |
| PhTox + Driver control | *Pre + Post-Gal4* >> + PhTox | 0.59 ± 0.04 | 0.5 ± .01 | 40.5 ± 1.0 | -67.0 ± 0.9 | 70.5 ± 4.3 *** | 150.6 ± 10.7 *** | 11 |
| *GluRIII* | *Pre + Post-Gal4* >> *UAS-GluRIII RNAi/+* | 0.64 ± 0.03 | 0.4 ± 0.0 | 37.5 ± 1.2 | -70.1 ± 1.2 | 58.9 ± 2.1 *** (vs. Driver control) | 110.6 ± 5.7 | 13 |
| *GluRIII* | *Pre + Post-Gal4* >> *UAS-GluRIII RNAi/+* + PhTox | 0.37 ± 0.01 | 0.2 ± 0.0 | 29.6 ± 1.0 | -65.8 ± 0.8 | 80.1 ± 3.5 *** (vs. *GluRIII* alone) | 133.2 ± 7.6 | 15 |
| wild type | | 0.78 ± 0.02 | 2.0 ± 0.1 | 36.6 ± 0.7 | -67.1 ± 0.6 | 48.3 ± 1.3 | 93.3 ± 3.0 | 57 |
| wild type | + PhTox | 0.53 ± 0.02 | 0.8 ± 0.1 | 33.3 ± 0.7 | -64.4 ± 0.3 | 65.1 ± 2.8 *** | 119.7 ± 6.3 *** | 24 |
| *Plc21C* (*GluRIII* cont) | *Pre + Post-Gal4* >> *GD11359* | 0.80 ± 0.04 | 1.3 ± 0.1 | 34.9 ± 1.4 | -67.4 ± 1.4 | 44.7 ± 2.8 | 83.7 ± 6.9 | 14 |
| *Plc21C* | *Pre + Post-Gal4* >> *GD11359/+* *GluRIII RNAi/+* | 0.55 ± 0.01 | 0.3 ± 0.0 | 29.6 ± 1.0 | -66.0 ± 0.7 | 53.9 ± 0.5 * (vs. *Plc21C RNAi*) | 89.4 ± 5.0 | 17 |
| *Plc21C* | *Pre + Post-Gal4* >> *GD11359/+* *GluRIII RNAi/+* + PhTox | 0.42 ± 0.01 | 0.2 ± 0.1 | 27.3 ± 1.7 | -64.9 ± 1.2 | 64.9 ± 4.1 * (vs. *Plc21C* + *GluRIII RNAi*) | 105.1 ± 10.4 | 11 |
| *Plc21C* (PhTox cont) | *Pre + Post-Gal4* >> *GD11359* | 0.80 ± 0.04 | 0.7 ± 0.1 | 31.6 ± 0.9 | -66.0 ± 0.9 | 40.6 ± 2.0 *** | 69.8 ± 6.5 | 14 |
| *Plc21C* | *Pre + Post-Gal4* >> *GD11359/+* + PhTox | 0.51 ± 0.01 | 0.3 ± 0.1 | 30.9 ± 1.7 | -66.5 ± 0.9 | 61.1 ± 3.7 *** | 106.8 ± 9.5 ** | 14 |

### Supplementary Table S2

| FIGURE 2 – SCREEN | | | | | | | | |
|---|---|---|---|---|---|---|---|---|
| **Condition** | **Genotype or Reagent** | **mEPSP (mV)** | **mEPSP freq. (Hz)** | **EPSP (mV)** | **$V_m$ (mV)** | **QC** | **NLSC QC** | **n** |
| wild type | $w^{1118}$ | 0.88 ± 0.02 | 3.3 ± 0.2 | 38.3 ± 0.7 | -68.2 ± 0.7 | 44.1 ± 1.0 | 87.6 ± 2.7 | 52 |
| *GluRIIA* | $GluRIIA^{SP16}$ | 0.46 ± 0.02 | 0.9 ± 0.1 | 29.2 ± 1.0 | -68.7 ± 0.8 | 66.0 ± 3.1 *** | 107.3 ± 6.2 *** | 24 |
| *GluRIII RNAi* | *Pre + Post-Gal4* | 0.81 ± 0.02 | 2.0 ± 0.7 | 37.5 ± 1.3 | -68.3 ± 1.0 | 46.7 ± 1.7 | 91.9 ± 5.0 | 22 |
| | *Pre + Post-Gal4 >> GluRIII* RNAi/+ | 0.57 ± 0.02 | 0.3 ± 0.0 | 34.1 ± 0.8 | -67.6 ± 1.0 | 60.5 ± 1.9 *** | 108.3 ± 3.9 * | 17 |
| *CG7611* | $CG7611^{EY07631}$ | 0.93 ± 0.03 | 3.3 ± 0.3 | 28.3 ± 1.3 | -67.3 ± 0.9 | 30.9 ± 1.7 | 49.6 ± 3.4 | 12 |
| | $GluRIIA^{SP16}$; $CG7611^{EY07631}$ | 0.45 ± 0.02 | 1.6 ± 0.2 | 23.6 ± 1.7 | -67.1 ± 1.0 | 53.1 ± 4.4 *** | 79.6 ± 8.8 ** | 13 |
| *Mipp2* | *Pre + Post-Gal4 >> TRiP.HMC03229* | 0.98 ± 0.04 | 1.1 ± 0.2 | 36.4 ± 1.5 | -69.8 ± 1.6 | 37.4 ± 1.5 | 69.5 ± 3.9 | 10 |
| | *Pre + Post-Gal4 >> TRiP.HMC03229 + GluRIII* RNAi | 0.65 ± 0.03 | 0.2 ± 0.0 | 33.7 ± 1.2 | -68.1 ± 1.2 | 52.5 ± 2.3 *** | 92.6 ± 4.7 ** | 9 |
| *Mipp2* | *Pre + Post-Gal4 >> 4317-R2* | 0.82 ± 0.03 | 0.8 ± 0.1 | 43.9 ± 1.7 | -71.4 ± 1.4 | 53.9 ± 2.3 | 118.9 ± 8.6 | 11 |
| | *Pre + Post-Gal4 >> 4317-R2 + GluRIII* RNAi | 0.60 ± 0.03 | 0.4 ± 0.0 | 37.0 ± 1.6 | -71.7 ± 1.1 | 62.0 ± 2.0 * | 113.9 ± 5.1 | 10 |
| *Mipp2* | $Mipp2^{KG01786}$ | 0.97 ± 0.04 | 1.7 ± 0.2 | 39.0 ± 1.1 | -69.8 ± 1.1 | 40.8 ± 1.7 | 81.1 ± 4.5 | 15 |
| | $Mipp2^{KG01786}$; $GluRIIA^{SP16}$ | 0.52 ± 0.03 | 1.1 ± 0.2 | 29.6 ± 1.7 | -71.7 ± 1.1 | 57.2 ± 3.4 *** | 92.4 ± 6.9 | 16 |
| *slowpoke* | $slo^4$ | 0.83 ± 0.08 | 5.0 ± 0.1 | 36.3 ± 1.6 | -71.3 ± 1.0 | 47.7 ± 2.8 | 87.0 ±5.6 | 18 |
| | $GluRIIA^{SP16}$; $slo^4$ | 0.42 ± 0.02 | 1.7 ± 0.1 | 26.0 ± 1.7 | -70.3 ± 1.0 | 64.1 ± 5.2 * | 100.4 ± 9.4 | 22 |
| *slowpoke* | $slo^4/slo^1$ | 0.93 ± 0.05 | 3.7 ± 0.4 | 35.9 ± 1.2 | -69.9 ± 0.9 | 40.1 ± 2.2 | 73.9 ± 4.5 | 19 |
| | $GluRIIA^{SP16}$; $slo^4/slo^1$ | 0.38 ± 0.01 | 1.4 ± 0.2 | 25.4 ± 1.6 | -70.8 ± 0.9 | 67.5 ± 4.0 *** | 102.6 ± 8.7 *** | 18 |
| *Gβ13F* | $G\beta13F^{KG08410}$ | 0.97 ± 0.06 | 1.4 ± 0.2 | 26.9 ± 1.9 | -63.2 ± 0.4 | 28.1 ± 2.3 | 45.6 ± 4.9 | 8 |
| | $GluRIIA^{SP16}$; $G\beta13F^{KG08410}$ | 0.52 ± 0.01 | 2.6 ± 0.6 | 31.8 ± 1.5 | -66.2 ± 0.1 | 61.1 ± 2.6 *** | 106.0 ± 7.2 *** | 7 |
| *Gβ76C* | $G\beta76C^1$ | 0.99 ± 0.05 | 1.0 ± 0.1 | 39.6 ± 0.9 | -69.8 ± 1.1 | 40.6 ± 1.9 | 80.9 ± 4.0 | 7 |
| | $GluRIIA^{SP16}$; $G\beta76C^1$ | 0.50 ± 0.02 | 0.7 ± 0.1 | 29.6 ± 2.0 | -71.9 ± 2.6 | 59.4 ± 2.6 *** | 93.9 ± 6.5 | 8 |

| Condition | Genotype or Reagent | mEPSP (mV) | mEPSP freq. (Hz) | EPSP (mV) | $V_m$ (mV) | QC | NLSC QC | n |
|---|---|---|---|---|---|---|---|---|
| *Gβ76C* | *Pre + Post-Gal4 >> GD 13785* | 0.73 ± 0.04 | 0.4 ± 0.0 | 41.1 ± 1.4 | -70.3 ± 0.9 | 57.9 ± 3.3 | 120.1 ± 7.8 | 13 |
| | *Pre + Post-Gal4 >> GD 13785 GluRIII* RNAi | 0.54 ± 0.02 | 0.3 ± 0.0 | 34.5 ± 1.4 | -69.3 ± 1.7 | 64.7 ± 0.7 | 115.6 ± 2.6 | 16 |
| *TkR86C* | *Pre + Post-Gal4 >> TRiP.JF02160* | 1.00 ± 0.06 | 0.4 ± 0.1 | 36.7 ± 2.0 | -66.7 ± 0.8 | 38.5 ± 3.6 | 77.5 ± 9.8 | 10 |
| | *Pre + Post-Gal4 >> GluRIII* RNAi/ *TRiP.JF02160* | 0.78 ± 0.02 | 0.3 ± 0.0 | 41.2 ± 1.1 | -66.9 ± 1.5 | 52.6 ± 2.0 ** | 113.9 ± 5.7 ** | 8 |
| *mAChR-A* | *Pre + Post-Gal4 >> TRiP.HMC02343* | 0.97 ± 0.03 | 1.1 ± 0.2 | 37.1 ± 1.1 | -66.3 ± 0.6 | 38.7 ± 1.5 | 76.2 ± 4.2 | 14 |
| | *Pre + Post-Gal4 >> TRiP.HMC02343 + GluRIII* RNAi | 0.71 ± 0.03 | 0.1 ± 0.0 | 34.1 ± 1.5 | -66.7 ± 0.8 | 48.5 ± 1.5 *** | 88.3 ± 4.3 * | 13 |
| *GABA-B-R1* | *Pre + Post-Gal4 >> TRiP.HMC03388* | 1.00 ± 0.05 | 0.7 ± 0.1 | 39.6 ± 1.4 | -69.8 ± 0.7 | 40.1 ± 1.5 | 80.2 ± 3.9 | 7 |
| | *Pre + Post-Gal4 >> TRiP.HMC03388 + GluRIII* RNAi | 0.78 ± 0.02 | 0.3 ± 0.0 | 35.1 ± 0.8 | -66.5 ± 1.2 | 45.1 ± 1.4 * | 83.5 ± 3.1 | 8 |
| *PK2-R2* | *Pre + Post-Gal4 >> TRiP.JF03209* | 1.02 ± 0.05 | 1.6 ± 0.5 | 36.0 ± 1.6 | -67.1 ± 1.0 | 35.7 ± 0.9 | 67.3 ± 2.4 | 8 |
| | *Pre + Post-Gal4 >> TRiP.JF03209 + GluRIII* RNAi | 0.80 ± 0.04 | 0.8 ± 0.1 | 35.2 ± 1.8 | -64.5 ± 0.8 | 44.2 ± 1.9 ** | 85.1 ± 5.6 * | 9 |
| *methuselah* | *Pre + Post-Gal4 >> TRiP.GL01060* | 0.81 ± 0.03 | 1.1 ± 0.2 | 37.5 ± 1.5 | -68.2 ± 1.1 | 46.9 ± 2.0 | 91.7 ± 5.5 | 15 |
| | *Pre + Post-Gal4 >> TRiP.GL01060 + GluRIII* RNAi | 0.66 ± 0.03 | 0.4 ± 0.1 | 36.7 ± 1.3 | -70.5 ± 1.1 | 56.4 ± 1.9 ** | 105.3 ± 5.2 | 13 |
| *AdoR* | *Pre + Post-Gal4 >> TRiP.JF02687* | 0.89 ± 0.05 | 1.5 ± 0.2 | 33.1 ± 1.7 | -69.3 ± 1.3 | 37.8 ± 2.5 | 65.8 ± 5.7 | 8 |
| | *Pre + Post-Gal4 >> TRiP.JF02687 + GluRIII* RNAi | 0.69 ± 0.03 | 0.6 ± 0.1 | 32.5 ± 1.9 | -67.6 ± 1.8 | 46.9 ± 2.4 * | 81.7 ± 5.9 | 9 |
| *PKC* | *Pre + Post-Gal4 >> UAS-PKCi.B* | 0.79 ± 0.04 | 1.0 ± 0.2 | 35.7 ± 1.7 | -70.3 ± 1.5 | 45.9 ± 2.1 | 83.6 ± 5.2 | 14 |
| | *Pre + Post-Gal4 >> UAS-PKCi.B + GluRIII* RNAi | 0.62 ± 0.03 | 0.2 ± 0.0 | 33.5 ± 1.0 | -66.7 ± 1.2 | 55.4 ± 3.1 * | 100.8 ± 8.0 | 14 |
| *unc-13* | *Pre + Post-Gal4 >> KK109346* | 1.12 ± 0.04 | 2.6 ± 0.2 | 41.6 ± 1.7 | -73.0 ± 1.4 | 37.5 ± 1.6 | 77.0 ± 4.7 | 15 |
| | *Pre + Post-Gal4 >> KK109346 + GluRIII* RNAi | 0.88 ± 0.03 | 0.8 ± 0.1 | 39.8 ± 1.2 | -68.6 ± 0.8 | 45.4 ± 1.1 *** | 92.7 ± 3.6 * | 15 |

| Condition | Genotype or Reagent | mEPSP (mV) | mEPSP freq. (Hz) | EPSP (mV) | $V_m$ (mV) | QC | NLSC QC | n |
|---|---|---|---|---|---|---|---|---|
| *iav* | *Pre + Post-Gal4 >> TRiP.JF01904* | 0.89 ± 0.03 | 0.9 ± 0.1 | 35.9 ± 1.0 | -70.7 ± 1.2 | 40.6 ± 1.4 | 73.8 ± 3.3 | 14 |
| | *Pre + Post-Gal4 >> TRiP.JF01904 + GluRIII* RNAi | 0.70 ± 0.16 | 0.4 ± 0.1 | 32.1 ± 1.2 | -67.7 ± 1.0 | 45.6 ± 1.6 * | 78.7 ± 4.1 | 14 |
| *Slip1* | *Pre + Post-Gal4 >> TRiP.HMC03268* | 0.70 ± 0.03 | 0.9 ± 0.1 | 31.2 ± 1.6 | -65.7 ± 0.6 | 45.2 ± 3.2 | 79.2 ± 8.0 | 11 |
| | *Pre + Post-Gal4 >> TRiP.HMC03268 + GluRIII* RNAi | 0.56 ± 0.03 | 0.3 ± 0.0 | 33.1 ± 1.9 | -67.7 ± 1.0 | 59.6 ± 3.1 ** | 105.7 ± 8.3 * | 9 |
| *mGluR* | *$mGluR^{MI02169}$* | 1.01 ± 0.05 | 7.0 ± 0.8 | 36.0 ± 1.1 | -64.3 ± 0.7 | 36.0 ± 2.3 | 70.5 ± 5.9 | 6 |
| | *$GluRIIA^{SP16}$;;$mGluR^{MI02169}$* | 0.45 ± 0.01 | 1.7 ± 0.9 | 28.7 ± 1.6 | -63.0 ± 0.7 | 64.4 ± 4.3 *** | 108.8 ± 11.7 | 7 |
| *Pka-R2* | *Pre + Post-Gal4 >> TRiP.JF02759* | 0.79 ± 0.04 | 1.0 ± 0.1 | 35.0 ± 1.4 | -64.9 ± 0.5 | 44.7 ± 1.7 | 84.5 ± 4.4 | 8 |
| | *Pre + Post-Gal4 >> TRiP.JF02759* | 0.58 ± 0.03 | 2.7 ± 2.4 | 31.4 ± 1.9 | -64.8 ± 1.5 | 54.3 ± 2.6 ** | 94.6 ± 6.9 | 7 |
| *Gβ5* | *Pre + Post-Gal4 >> TRiP.JF02941* | 1.15 ± 0.07 | 1.6 ± 0.2 | 39.4 ± 1.7 | -70.6 ± 2.2 | 35.0 ± 1.8 | 69.0 ± 4.0 | 13 |
| | *Pre + Post-Gal4 >> TRiP.JF02941 GluRIII RNAi* | 0.80 ± 0.06 | 0.3 ± 0.1 | 39.9 ± 1.8 | -69.5 ± 1.1 | 50.6 ± 1.9 *** | 102.0 ± 4.3 *** | 11 |
| *CaMKII* (presynaptic) | *Pre-Gal4 >> UAS-CaMKII-Ala* | 0.90 ± 0.03 | 2.0 ± 0.2 | 38.3 ± 1.5 | -69.3 ± 1.0 | 43.2 ± 1.9 | 85.0 ± 5.4 | 15 |
| | *Pre-Gal4 >> UAS-CaMKII-Ala, GluR/GluR* | 0.51 ± 0.02 | 0.8 ± 0.1 | 28.3 ± 1.7 | -67.4 ± 1.1 | 56.6 ± 3.5 ** | 91.1 ± 7.2 | 11 |
| | *Pre-Gal4 >> UAS-CaMKII-Ala + PhTox* | 0.53 ± 0.02 | 1.2 ± 0.1 | 31.9 ± 2.6 | -66.5 ± 1.0 | 62.1 ± 6.3 *** | 113.66 ± 16.33 | 9 |

| Condition | Genotype or Reagent | mEPSP (mV) | mEPSP freq. (Hz) | EPSP (mV) | $V_m$ (mV) | QC | NLSC QC | n |
|---|---|---|---|---|---|---|---|---|
| *UAS-IP$_3$-sponge.m49* | *Pre + Post-Gal4 >> UAS-IP3-sponge.m49* | 0.89 ± 0.04 | 1.3 ± .01 | 43.1 ± 1.7 | -69.3 ± 1.3 | 49.1 ± 2.0 | 111.1 ± 8.7 | 13 |
| | *Pre + Post-Gal4 >> UAS-IP$_3$-sponge.m49 + GluRIII* RNAi | 0.55 ± 0.03 | 0.3 ± 0.1 | 28.5 ± 1.4 | -64.4 ± 0.5 | 52.4 ± 2.4 | 86.4 ± 5.2 * | 14 |
| *UAS-IP$_3$-sponge.m30* | *Pre + Post-Gal4 >> UAS-IP$_3$-sponge.m30* | 0.81 ± 0.04 | 1.4 ± 0.2 | 43.8 ± 1.4 | -69.4 ± 1.6 | 55.1 ± 1.8 | 123.3 ± 4.6 | 16 |
| | *Pre + Post-Gal4 >> UAS-IP$_3$-sponge.m30 + GluRIII* RNAi | 0.63 ± 0.02 | 0.3 ± 0.0 | 34.6 ± 1.1 | -67.2 ± 0.9 | 55.5 ± 2.3 | 102.0 ± 5.4 ** (down) | 15 |
| *UAS-IP$_3$-sponge.m49* (PhTox) | *Pre + Post-Gal4 >> UAS-IP$_3$-sponge.m49* (PhTox Cont) | 0.90 ± 0.03 | 1.6 ± 0.2 | 40.1 ± 1.3 | -67.4 ± 1.0 | 45.2 ± 2.4 | 96.5 ± 7.8 | 13 |
| | *Pre + Post-Gal4 >> UAS-IP$_3$-sponge.m49* PhTox | 0.54 ± 0.02 | 1.0 ± 0.1 | 38.6 ± 2.4 | -65.3 ± 0.9 | 73.1 ± 5.0 *** | 163.2 ± 17.6 ** | 15 |
| *UAS-IP$_3$-sponge.m49* | *Pre + Post-Gal4 >> UAS-IP$_3$-sponge.m49 + GluRIII* RNAi | 0.66 ± 0.03 | 0.4 ± 0.1 | 31.7 ± 1.2 | -64.9 ± 0.7 | 48.9 ± 1.8 | 86.4 ± 4.7 | 16 |
| | *Pre + Post-Gal4 >> UAS-IP$_3$-sponge.m49 + GluRIII* RNAi PhTox | 0.40 ± 0.02 | 0.2 ± 0.0 | 27.5 ± 1.8 | -64 ± 0.7 | 69.9 ± 4.9 *** | 116.1 ± 12.1 * | 14 |

**Supplementary Table S3**

| FIGURE 4 | | | | | | | | |
|---|---|---|---|---|---|---|---|---|
| **Condition** | **Genotype or Reagent** | **mEPSP (mV)** | **mEPSP freq. (Hz)** | **EPSP (mV)** | **$V_m$ (mV)** | **QC** | **NLSC QC** | **n** |
| wild type | 5 µM Xestospongin C | 0.85 ± 0.04 | 3.1 ± 0.3 | 38.8 ± 1.1 | -66.4 ± 0.9 | 46.3 ± 2.3 | 95.2 ±0.5 | 13 |
| *GluRIIA$^{SP16}$* | 5 µM Xestospongin C | 0.46 ± 0.03 | 0.8 ± 0.1 | 28.5 ± 1.9 | -68.2 ± 1.5 | 64.0 ± 4.3 * | 104.3 ± 9.8 | 15 |
| wild type | 20 µM Xestospongin C | 1.00 ± 0.04 | 6.1 ± 0.4 | 37.3 ± 0.8 | -64.1 ± 0.6 | 37.7 ± 0.9 | 76.4 ± 2.5 | 14 |
| wild type | 20 µM PhTox + 20 µM Xestospongin C | 0.56 ± 0.01 | 1.5 ± 0.2 | 34.2 ± 1.0 | -64.6 ± 0.8 | 61.8 ± 1.6 *** | 115.9 ± 4.9 *** | 19 |
| *GluRIIA$^{SP16}$* | 20 µM Xestospongin C | 0.54 ± 0.02 | 1.3 ± 0.2 | 21.4 ± 0.9 | -64.1 ± 0.1 | 40.4 ± 2.1 | 57.4 ± 3.5 *** (down) | 13 |
| wild type | DMSO (2% v/v) | 0.85 ± 0.04 | 5.4 ± 0.5 | 35.8 ± 1.2 | -64.9 ± 0.8 | 42.8 ± 1.8 | 83.1 ± 4.5 | 12 |
| *GluRIIA$^{SP16}$* | DMSO (2% v/v) | 0.46 ± 0.02 | 1.21 ± 0.16 | 26.1 ± 1.1 | -65.2 ± 0.7 | 57.5 ± 3.4 *** | 89.5 ± 7.0 | 11 |
| wild type | PhTox + DMSO (2% v/v) | 0.56 ± 0.02 | 1.9 ± 0.2 | 35.7 ± 1.0 | -67.0 ± 1.1 | 64.5 ± 2.6 *** | 121.2 ± 5.7 *** | 15 |

**Supplementary Table S4**

| FIGURE 5 | | | | | | | | |
|---|---|---|---|---|---|---|---|---|
| **Condition** | **Genotype or Reagent** | **mEPSP (mV)** | **mEPSP freq. (Hz)** | **EPSP (mV)** | **$V_m$ (mV)** | **QC** | **NLSC QC** | **n** |
| wild type (GluR cont) | 1 µM 2-APB | 0.80 ± 0.02 | 3.6 ± 0.3 | 35.9 ± 1.3 | -68.0 ± 1.1 | 45.0 ± 1.3 | 85.2 ± 4.3 | 15 |
| *GluRIIA$^{SP16}$* | 1 µM 2-APB | 0.53 ± 0.02 | 0.9 ± 0.1 | 23.4 ± 1.1 | -67.2 ± 0.8 | 45.3 ± 2.6 | 66.4 ± 5.1 ** (down) | 14 |
| wild type (PhTox cont) | 1 µM 2-APB | 0.75 ± 0.02 | 2.9 ± 0.2 | 39.9 ± 1.2 | -66.0 ± 0.9 | 53.5 ± 1.8 | 114.2 ± 6.0 | 11 |
| wild type | 1 µM 2-APB 20 µM PhTox | 0.48 ± 0.04 | 1.6 ± 0.3 | 35.6 ± 1.8 | -63.2 ± 0.5 | 76.9 ± 4.0 *** | 152.3 ± 9.5 ** | 12 |
| wild type | 10 µM 2-APB | 0.83 ± 0.03 | 4.7 ± 0.9 | 43.3 ± 0.8 | -65.0 ± 0.7 | 53.3 ± 2.5 | 128.3 ± 8.1 | 15 |
| *GluRIIA$^{SP16}$* | 10 µM 2-APB | 0.44 ± 0.02 | 0.7 ± 0.1 | 23.9 ± 1.0 | -64.9 ± 0.6 | 54.9 ± 2.9 | 82.2 ± 5.8 *** | 14 |

### Supplementary Table S5

| FIGURE 6 | | | | | | | | |
|---|---|---|---|---|---|---|---|---|
| **Condition** | **Genotype or Reagent** | **mEPSP (mV)** | **mEPSP freq. (Hz)** | **EPSP (mV)** | **$V_m$ (mV)** | **QC** | **NLSC QC** | **n** |
| wild type | +100 µM Ryanodine | 0.78 ± 0.02 | 3.7 ± 0.3 | 34.6 ± 1.1 | -66.4 ± 0.9 | 45.0 ± 2.2 | 84.1 ± 5.4 | 17 |
| *GluRIIA*$^{SP16}$ | +100 µM Ryanodine | 0.50 ± 0.01 | 1.3 ± 0.1 | 25.4 ± 1.1 | -66.2 ± 0.8 | 51.5 ± 2.4 | 78.4 ± 4.8 | 15 |
| wild type | +10 µM dantrolene | 0.81 ± 0.04 | 3.2 ± 0.2 | 29.2 ± 1.6 | -64.3 ± 0.5 | 37.3 ± 2.4 | 65.0 ± 5.1 | 25 |
| *GluRIIA*$^{SP16}$ | +10 µM dantrolene | 0.46 ± 0.01 | 0.9 ± 0.1 | 15.9 ± 1.3 | -64.8 ± 0.6 | 34.7 ± 2.9 | 46.5 ± 4.7 ** | 27 |
| wild type | +10 µM dantrolene | 0.91 ± 0.04 | 2.6 ± 0.2 | 34.8 ± 1.6 | -64.2 ± 0.7 | 39.4 ± 2.6 | 76.5 ± 6.5 | 13 |
| wild type | +PhTox +10 µM dantrolene | 0.53 ± 0.02 | 1.4 ± 0.2 | 34.4 ± 1.3 | -64.7 ± 0.9 | 66.1 ± 4.8 *** | 126.3 ± 13.7 ** | 10 |
| *GluRIII* RNAi | *Pre + Post-Gal4 >> GluRIII* RNAi/+ 10 µM Dantrolene | 0.06 ± 0.02 | 0.5 ± 0.1 | 37.1 ± 1.9 | -66.7 ± 1.7 | 66.9 ± 4.2 | 131.3 ± 11.8 | 8 |
| *GluRIII* RNAi | *Pre + Post-Gal4 >>* GluRIII RNAi +20 µM PhTox +10 µM dantrolene | 0.39 ± 0.03 | 0.4 ± 0.1 | 36.2 ± 1.5 | -67.4 ± 1.2 | 96.5 ± 6.0 ** | 183.8 ± 13.7 * | 11 |

**Supplementary Table S6**

| FIGURE 7 | | | | | | | | |
|---|---|---|---|---|---|---|---|---|
| **Condition** | **Genotype or Reagent** | **mEPSP (mV)** | **mEPSP freq. (Hz)** | **EPSP (mV)** | **$V_m$ (mV)** | **QC** | **NLSC QC** | **n** |
| Driver Control | *Pre + Post-Gal4 >>* +10 µM Dantrolene | 0.73 ± 0.03 | 0.9 ± 0.1 | 39.9 ± 1.8 | -67.6 ± 0.9 | 55.5 ± 2.7 | 118.2 ± 9.5 | 14 |
| *GluRIII* RNAi | *Pre + Post-Gal4 >> GluRIII* RNAi/+ 10 µM Dantrolene | 0.61 ± 0.02 | 0.2 ± 0.0 | 31.5 ± 1.0 | -65.5 ± 0.9 | 52.6 ± 2.2 | 90.8 ± 4.7 | 14 |
| Driver Control *$IP_3$ sponge* | *Pre + Post-Gal4 >> UAS-$IP_3$-sponge.m49* 10 µM Dantrolene | 0.93 ± 0.03 | 1.6 ± 0.1 | 41.2 ± 1.7 | -67.6 ± 1.4 | 44.5 ± 1.5 | 96.1 ± 5.8 | 12 |
| *GluRIII* RNAi *$IP_3$ sponge* | *Pre + Post-Gal4 >> GluRIII* RNAi/*UAS-$IP_3$-sponge.m49* 10 µM Dantrolene | 0.68 ± 0.05 | 0.4 ± 0.1 | 33.3 ± 1.8 | -66.0 ± 1.1 | 50.3 ± 2.1 | 90.5 ± 4.3 | 13 |
| wild type | | 0.78 ± 0.04 | 1.8 ± 0.2 | 33.7 ± 0.9 | -64.7 ± 0.5 | 44.6 ± 1.8 | 82.4 ± 4.0 | 20 |
| wild type | + PhTox | 0.65 ± 0.03 | 1.4 ± 0.2 | 36.5 ± 1.0 | -65.0 ± 1.1 | 57.2 ± 3.5 ** | 112.0 ± 7.8 ** | 12 |
| Driver control | *Pre + Post-Gal4* | 0.68 ± 0.04 | 2.0 ± 0.8 | 40.1 ± 1.6 | -66.4 ± 1.0 | 61.1 ± 4.0 | 132.9 ± 12.5 | 13 |
| Driver control | *Pre + Post-Gal4* PhTox CNS intact | 0.57 ± 0.03 | 0.7 ± 0.8 | 39.8 ± 0.8 | -66.9 ± 1.3 | 72.5 ± 4.4 | 152.7 ± 10.3 | 12 |
| Driver control | *Pre + Post-Gal4* PhTox CNS excised | 0.52 ± 0.02 | 0.7 ± 0.1 | 37.8 ± 1.4 | -66.0 ± 1.3 | 72.9 ± 3.6 * | 149.7 ± 12.6 | 12 |
| *UAS-$IP_3$-sponge.m49* | *Pre + Post-Gal4 >> UAS-$IP_3$-sponge.m49* | 0.83 ± 0.04 | 1.3 ± 0.1 | 40.4 ± 1.2 | -64.9 ± 0.8 | 50.1 ± 202 | 111.2 ± 6.8 | 15 |
| *UAS-$IP_3$-sponge.m49* | *Pre + Post-Gal4 >> UAS-$IP_3$-sponge.m49* PhTox | 0.56 ± 0.02 | 1.0 ± 0.1 | 40.4 ± 1.8 | -65.6 ± 1.0 | 72.8 ± 3.8 *** | 161.9 ± 12.8 ** | 13 |
| *UAS-$IP_3$-sponge.m49* | *Pre + Post-Gal4 >> UAS-$IP_3$-sponge.m49* PhTox | 0.56 ± 0.04 | 0.7 ± 0.1 | 34.4 ± 1.6 | -64.9 ± 0.9 | 65.0 ± 5.5 * | 125.6 ± 14.7 | 13 |

**Supplementary Table S7**

| FIGURE 8 | | | | | | | | |
|---|---|---|---|---|---|---|---|---|
| **Condition** | **Genotype or Reagent** | **mEPSP (mV)** | **mEPSP freq. (Hz)** | **EPSP (mV)** | **$V_m$ (mV)** | **QC** | **NLSC QC** | **n** |
| Driver Control (Neuron) | *Pre-Gal4* | 0.95 ± 0.04 | 2.6 ± 0.2 | 38.7 ± 1.1 | -66.5 ± 1.0 | 41.9 ± 2.3 | 86.2 ± 5.8 | 17 |
| | *Pre-Gal4 $GluRIIA^{SP16}$* | 0.49 ± 0.01 | 1.3 ± 0.2 | 29.5 ± 1.2 | -65.1 ± 0.7 | 60.3 ± 2.2 *** | 101.2 ± 6.1 | 15 |
| *$IP_3$-sponge* | *Pre-Gal4 >> UAS-$IP_3$-sponge.m49* | 1.09 ± 0.04 | 3.5 ± 0.0 | 40.8 ± 1.5 | -69.9 ± 1.3 | 37.6 ± 1.3 | 77.9 ± 3.7 | 16 |
| *GluRIIA $IP_3$-sponge* | *Pre-Gal4 >> UAS-$IP_3$-sponge.m49 $GluRIIA^{SP16}$* | 0.47 ± 0.01 | 0.6 ± 0.1 | 24.3 ± 2.1 | -68.4 ± 1.1 | 51.6 ± 4.0 ** | 79.2 ± 9.4 | 15 |
| Driver Control (Muscle) | Post-Gal4 | 0.76 ± 0.0 | 2.8 ± 0.2 | 38.7 ± 1.0 | -67.6 ± 1.0 | 51.6 ± 2.1 | 103.6 ± 5.3 | 14 |
| | Post-Gal4 *$GluRIIA^{SP16}$* | 0.41 ± 0.01 | 0.7 ± 0.1 | 26.8 ± 1.0 | -66.2 ± 0.8 | 66.7 ± 3.0 *** | 105.4 ± 6.6 | 14 |
| *$IP_3$-sponge* | *Post-Gal4 >> UAS-$IP_3$-sponge.m49* | 0.85 ± 0.03 | 2.1 ± 0.2 | 38.3 ± 1.1 | -65.7 ± 0.7 | 45.4 ± 1.5 | 93.6 ± 4.8 | 15 |
| *GluRIIA $IP_3$-sponge* | *Post-Gal4 >> UAS-$IP_3$-sponge.m49 $GluRIIA^{SP16}$* | 0.51 ± 0.03 | 1.4 ± 0.2 | 27.8 ± 1.3 | -65.3 ± 0.9 | 56.2 ± 0.7 ** | 92.1 ± 7.9 | 16 |
